# High Throughput Information Extraction of Printed Specimen Labels from Large-Scale Digitization of Entomological Collections using a Semi-Automated Pipeline

**DOI:** 10.1101/2025.07.09.663724

**Authors:** Margot Belot, Joël Tuberosa, Leonardo Preuss, Olha Svezhentseva, Magdalena Claessen, Christian Bölling, Franziska Schuster, Théo Léger

**Affiliations:** Museum für Naturkunde, Leibniz-Institut für Evolutions- und Biodiversitätsforschung, Invalidenstraße 43, 10115 Berlin, Germany

**Keywords:** artificial intelligence, biodiversity informatics, CNN, insect digitization, label extraction, museum collections, OCR, text clustering

## Abstract

1. Natural history museums curate billions of insect specimens, forming a vast but underutilized resource for biodiversity research. While digitization efforts have increased the availability of high-resolution specimen images, extracting metadata from labels remains a major bottleneck, often requiring manual transcription.
2. We developed a semi-automated pipeline, ELIE (Entomological Label Information Extraction), which combines computer vision, convolutional neural networks (CNNs), optical character recognition (OCR), and clustering algorithms to streamline label data extraction. Our pipeline operates in three stages: (1) label detection and classification (printed vs. handwritten), (2) OCR-based text extraction from printed labels using Tesseract or Google Vision, and (3) clustering of extracted text for human validation of outliers.
3. Benchmarking on diverse datasets from multiple museum collections showed that our approach successfully extracted and clustered up to 98% of printed labels, significantly reducing manual effort. The pipeline improves efficiency in digitization workflows while maintaining high accuracy in label data capture.
4. Our approach demonstrates the potential of integrating AI-driven methods with human validation to accelerate specimen digitization. By reducing manual transcription workload and enabling scalable extraction of insect label metadata, it unlocks biodiversity data for research in ecology, systematics, and conservation globally.

## Introduction

Insects, with over a million described species, represent the largest group of living organisms and comprise an estimated 80% of the world’s undescribed biodiversity (Stork, 2017). They play vital ecological roles but are increasingly threatened by habitat loss, agriculture, and climate change (Halsch et al., 2021; Platts et al., 2019; Raven & Wagner, 2021). Over the past 300 years, entomologists have collected and preserved insect specimens, resulting in more than half a billion records housed in museum collections worldwide (Short et al., 2018). Entomological collections serve as the basis of systematic classification by storing the valuable type specimens (which link a taxon description to a physical specimen) and represent a reservoir of species yet to be named (Fontaine et al., 2012). Entomological collections further provide a unique opportunity to explore species distribution and abundance (Gotelli et al., 2021; Ponder et al., 2001), ecological networks (e.g., de Araújo Romeiro et al. 2023; Pérez-Lachaud et al., 2017; Rakosy et al., 2022), temporal variations in morphological traits (e.g., Arce et al. 2022; Bartomeus et al., 2018; Brooks et al., 2014; Freedman et al., 2020; Wilson et al., 2022), as well as in genomic traits (e.g., Cridland et al. 2018; Short et al. 2018). They also represent an invaluable source for investigating the impact of climate change (Kharouba et al., 2018).

Digitizing entomological collections holds the promise of unlocking data to study insect evolution over time and across various geographic scales. Ensuring these collections are accessible is particularly important for preserving material from developing countries, enabling them to retain access to their biological heritage while also fostering global research, conservation, and educational efforts (Poske, 2024). Extensive digitization initiatives have emerged in recent years, including taxon- or region-based projects aimed at building comprehensive databases focused on specific insect groups, such as AntWeb (California Academy of Sciences), LepNet (Lepidoptera of North America Network), and the BigBee project (USDA–ARS). Institution-led efforts further complement these projects by digitizing collections based on particular taxa or collection types, often guided by community interest. Many of the world’s largest natural history institutions have launched large-scale digitization projects to make their collections more accessible to the public. For instance, the Natural History Museum in London (NHM) recently digitized over 2.4 million insect specimens (Natural History Museum London, 2025). Similarly, the Smithsonian Institution has made significant progress, with digital records for nearly 600,000 insect specimens (Smithsonian, pers. comm.). The Museum für Naturkunde in Berlin (MfN) digitized 650,000 specimens between 2022 and 2023, employing an innovative conveyor belt system developed by Picturae (Leiden, Netherlands) that can capture images of up to 1,600 specimens daily (1300/day on average), and, in between, 26,000 and 32,000 specimens monthly (MfN, 2025).

In entomological collections, labels placed under pinned specimens bear most of the information about a specimen, with more than 85% of available specimen information currently residing on these labels or in physical ledgers (Walton et al., 2020a). These labels provide crucial data about the specimens, including the place and date of collection, taxonomic identification, and voucher linking to other objects such as collection catalogs, dissections, or DNA vouchers. Different methods have been proposed to accelerate the digitization of insect specimen labels, including angled imaging of the label or imaging with a smartphone (Price et al., 2018; Ahrens et al., 2025). While a system to extract and database data from specimen labels has been proposed for herbarium specimens (Kirchhoff et al., 2018; Takano et al., 2024), the extraction and processing of metadata information from insect specimens still heavily depend on human operators to this day. Manually transcribing label data, might it be by crowdsourcing, directly entering information into collection management systems, or outsourcing to external companies, is still the standard approach in museum institutions (Walton et al., 2020b). It has been estimated that 90% of the time spent on digitizing a specimen goes into labeling the specimen and capturing metadata (Blagoderov et al., 2012). However, this approach is slow, labor-intensive, and expensive, posing challenges for many institutions. Walton et al. (2020b) found that transcribing and extracting information from insect specimens’ labels took, on average, 5.77 minutes for handwritten labels and 4.80 minutes for printed labels. While OCR has been available for a couple of years now, institutions are still hesitant to use it due to the variable quality of the results (Walton et al., 2020b). Given the typically limited financial resources allocated to natural history collections, it is essential to develop workflows that leverage available technologies to process label information and reduce associated costs.

While manual transcription of entomological labels remains labor-intensive and time-consuming, automated approaches promise greater scalability, albeit with their challenges. OCR systems benefit from the abundance of high-quality training data available for printed text, enabling high accuracy with simpler models, such as CNNs. In contrast, handwritten text recognition (HTR) systems face significant challenges due to the variability and complexity of handwriting, necessitating more sophisticated models and extensive data generation techniques to acquire sufficient training data (Ingle et al., 2019). The performance of OCR on handwritten text varies depending on the quality of the text analyzed. According to a study by Ptucha et al. (2018), the Character Error Rate (CER) ranges from 2.5% to 10.1%, and the Word Error Rate (WER) ranges from 4.8% to 18.8% on a standardized dataset. The lower performance of OCR models on handwriting suggests a need for differential processing of the label images to optimize the results. Despite various concepts and proposed processes, as indicated by Owen et al. (2020), none have yet provided a practical, adaptable, and cost-effective solution for automating the extraction of entomological label information from extensive 2D imaging outputs.

To address these challenges, we hypothesized that (1) Separating printed from handwritten labels before OCR would improve overall text recognition accuracy; (2) Clustering OCR output based on textual similarity would allow for effective identification and deduplication of recurring labels; and (3) A modular, scalable pipeline addressing these components could significantly reduce manual transcription time without compromising data integrity.

The pipeline presented here, ELIE (Entomological Label Information Extraction), developed in Python 3, is optimized for interoperability and modularity (Fig. 1). Recognizing that many labels are either duplicates or share highly similar structure and content, with a significant number also containing printed text, the pipeline carries out eight actions: (1) A trained CNN detects and crops single-label 2D images from multi-label 2D images; (2) Empty labels are identified and filtered out; (3) A separate CNN distinguishes specimen identifier labels from other labels; (4) Another CNN classifies each remaining label as printed or handwritten; (5) Printed labels undergo preprocessing, including orientation correction; (6) Printed labels are processed with OCR, while handwritten ones are routed to either HTR systems or manual transcription; (7) OCR results are refined through Natural Language Processing (NLP)-based post-processing; (8) Finally, the content of all printed labels is clustered based on similarity, and the two most-distant labels are manually checked to validate these clusters. Many of these labels are either identical (e.g., collection data labels from the same batch) or differ only by serial numbers (e.g., DNA extraction or dissection labels).

**Figure 1.**
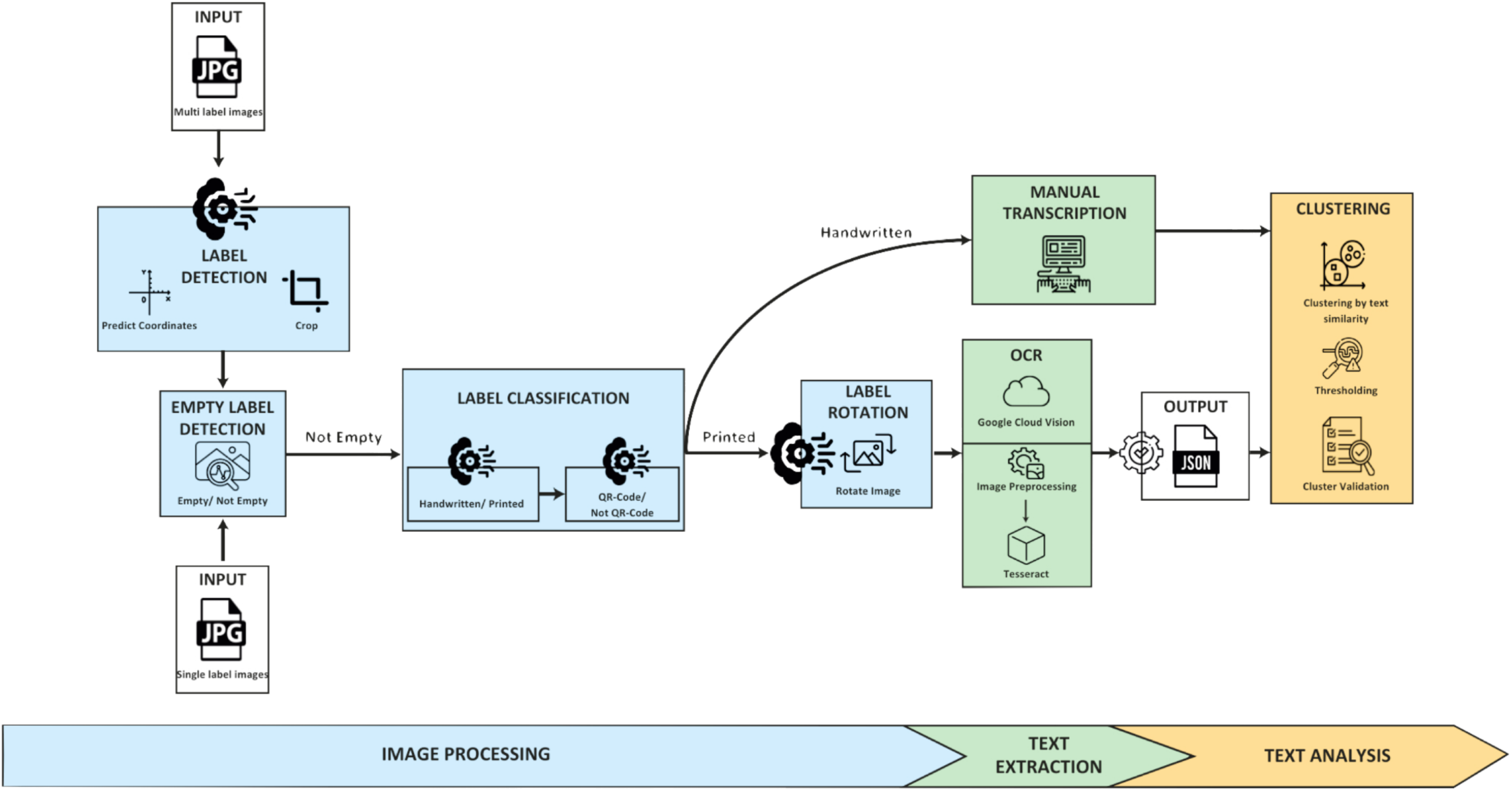
Overview of the ELIE (Entomological Label Information Extraction) pipeline. The flowchart illustrates the modular structure of the pipeline, which consists of three main stages: (1) image processing, (2) text extraction, and (3) text analysis. Each stage comprises independent modules, including label detection, empty label detection, classification (for printed vs. handwritten text and identifier detection), rotation, OCR processing, image post-processing, and label clustering. The modular design allows components to be activated selectively depending on dataset characteristics and digitization goals.

## Materials and Methods

### Data Collection

A total of 14,717 entomological label images were gathered to train, validate, and test the pipeline’s modules (Table 1). Multi-label images (MLI, fig. 2) from AntWeb, Bees & Bytes, LEPPHIL, and Picturae_MfN were used to support model training. Single-label images (SLI) from Picturae_MfN were used for classification and rotation tasks (Tables 2 and 3, in the Supporting Information). Three test datasets, from the MfN, the National Museum of Natural History (USNM), and the Museum of Comparative Zoology at Harvard University (MCZ), assessed pipeline performance (Table 4).

**Figure 2.**
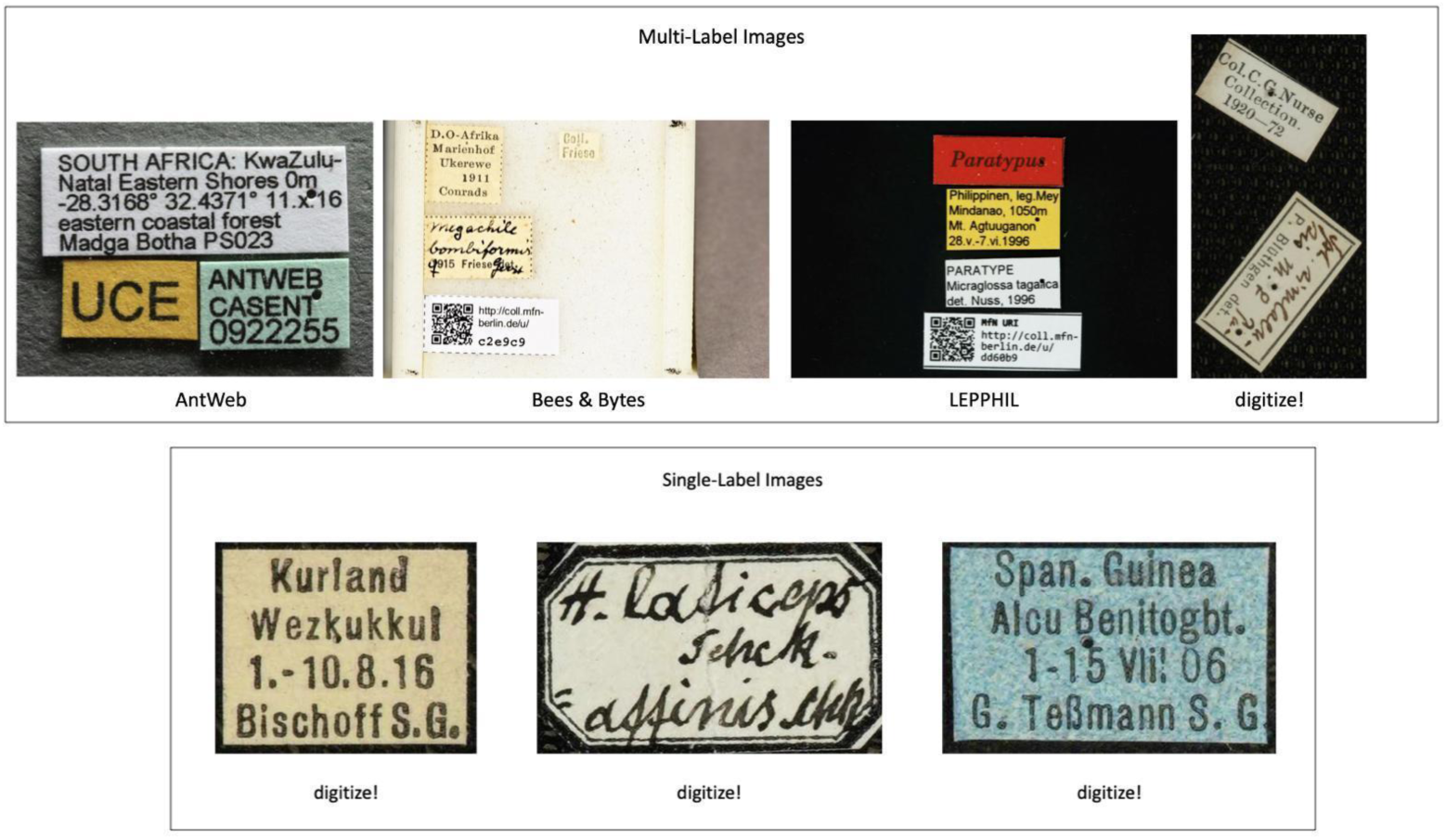
Examples of entomological specimen images used in the ELIE pipeline. Multi-label image (MLI) showing multiple labels beneath a pinned specimen. Single-label image (SLI) cropped from a multi-label image using the digitization system developed by Picturae (Leiden, Netherlands). Images are not to scale. Image credit: The MLI panel shows a specimen image from AntWeb (https://www.antweb.org/specimen/CASENT0919852), a photo by Michele Esposito. Used with permission, under a Creative Commons Attribution License (CC BY 4.0).

**Table 1.**
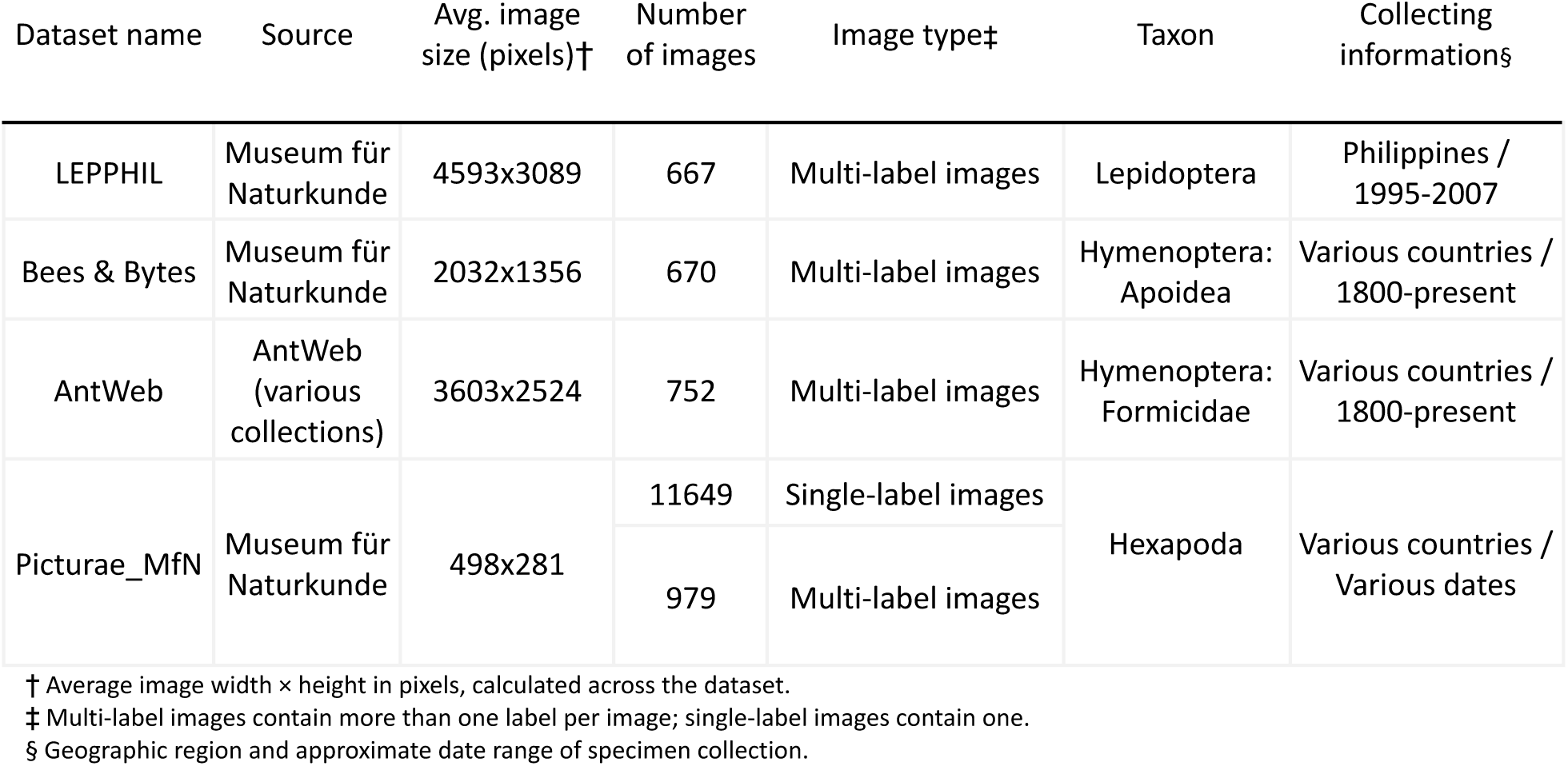
Overview of the entomological label datasets used for training, validation, and testing of the pipeline modules. All datasets include images of insect specimens collected from various locations and times.

| Dataset name | Source | Avg. image size (pixels)† | Number of images | Image type‡ | Taxon | Collecting informations§ |
| --- | --- | --- | --- | --- | --- | --- |
| LEPPHIL | Museum für Naturkunde | 4593x3089 | 667 | Multi-label images | Lepidoptera | Philippines / 1995-2007 |
| Bees & Bytes | Museum für Naturkunde | 2032x1356 | 670 | Multi-label images | Hymenoptera: Apoidea | Various countries / 1800-present |
| AntWeb | AntWeb (various collections) | 3603x2524 | 752 | Multi-label images | Hymenoptera: Formicidae | Various countries / 1800-present |
| Picturae_MfN | Museum für Naturkunde | 498x281 | 11649 | Single-label images | Hexapoda | Various countries / Various dates |
|  |  |  | 979 | Multi-label images |  |  |

† Average image width × height in pixels, calculated across the dataset.
‡ Multi-label images contain more than one label per image; single-label images contain one.
§ Geographic region and approximate date range of specimen collection.

**Table 4.** Overview of the entomological label datasets used for performance assessment of the pipeline. Each dataset includes labeled insect specimen images from major museum collections, varying in taxonomic focus, geographic origin, and image characteristics.

| Dataset name | Source | Avg. image size (pixels)† | Number of images | Image type‡ | Taxon | Collecting informations§ |
| --- | --- | --- | --- | --- | --- | --- |
| MfN_LEP_SEASIA | Museum für Naturkunde Berlin | 535.58x344.51 | 25844 | Single-label images | Lepidoptera: Pyraloidea | Southeast Asia / Various dates |
| MCZ_ENT_Boston | Museum of Comparative Zoology at Harvard University | 1000.00x932.89 | 1596 | Multi-label images | Hexapoda | North America / Various dates |
| USNM_COL_CAM | Smithsonian National Museum of Natural History | 3030x2080 | 912 | Multi-label images | Coleoptera: Buprestidae, Carabidae, Cerambycidae, Chrysomelida, Curculionidae, Scarabaeidae | South and Central America / Various dates |

† Average image width × height in pixels, calculated across the dataset.
‡ Multi-label images contain more than one label per image; single-label images contain one.
§ Geographic region and approximate date range of specimen collection.

### Stage 1: Image Processing

The ELIE label detection module extracts SLI from MLI using a pre-trained Faster R-CNN ResNet-50 FPN model (Bi, 2022). Trained on 3,058 annotated MLIs from LEPPHIL, Bees & Bytes, AntWeb, and Picturae_MfN, the model uses an 80-20-10 train-validation-test split (Raschka et al., 2019; Table 5, in the Supporting Information). Annotations were created in PASCAL VOC format with LabelImg, marking label boundaries and assigning a “label” class (PyPI, 2021; Fig. 3, in the Supporting Information). Performance is evaluated using Intersection over Union (IOU) scores, with accuracy defined as the proportion of predictions exceeding an IOU of 0.8 (Fig. 4.1).

**Figure 4.**
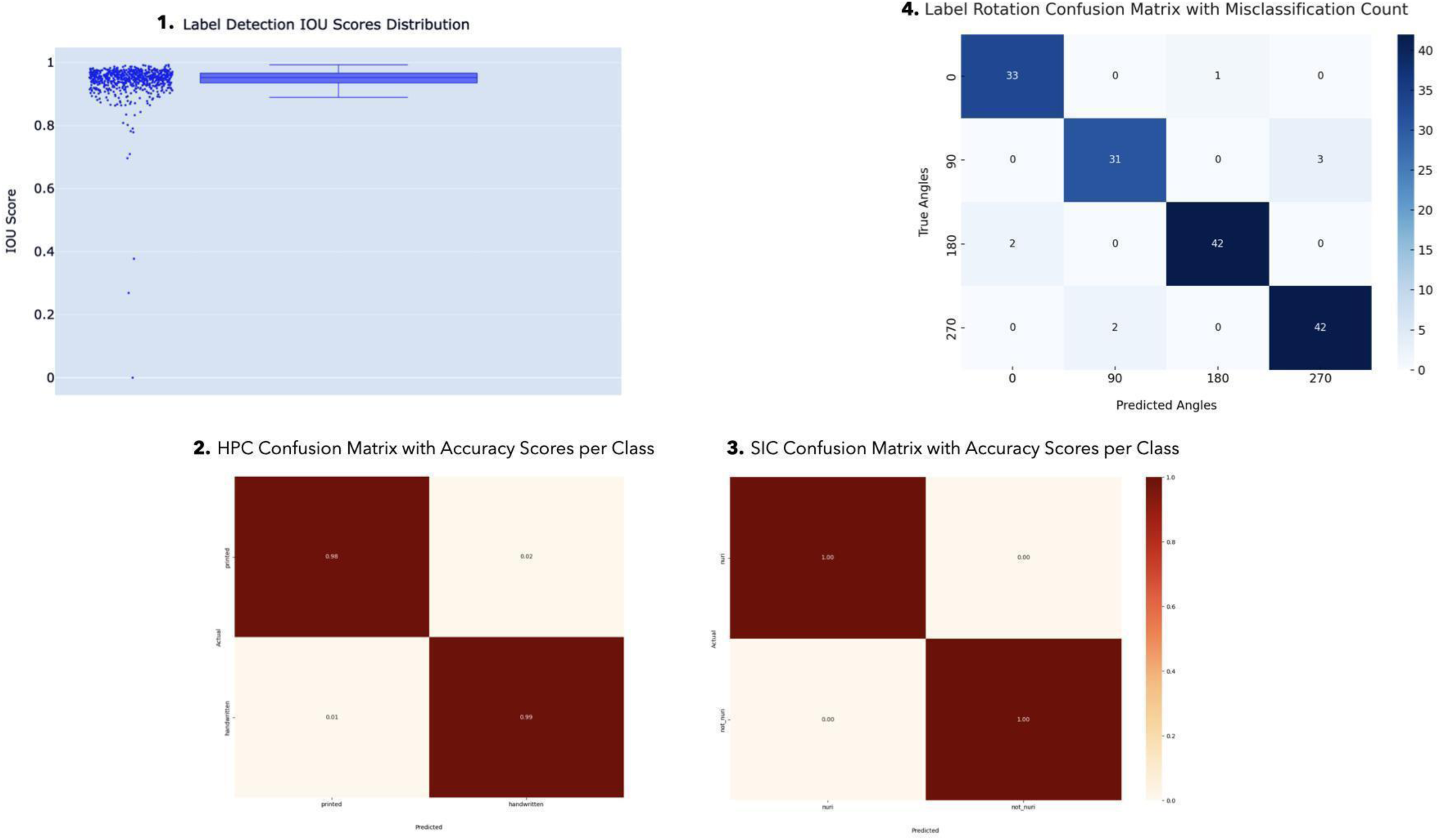
Evaluation of image processing modules in the ELIE pipeline. (4.1) Box plot of Intersection over Union (IOU) scores for the label detection module on the test dataset; each dot represents a predicted label. The horizontal line at IOU = 0.8 marks the accuracy threshold for correct detection. (4.2) Confusion matrix showing classification results of the Handwritten vs. Printed Classifier (HPC). (4.3) Confusion matrix for the Specimen Identifier Classifier (SIC), distinguishing identifier labels from all other label types. (4.4) Confusion matrix for the label rotation module, showing predicted versus ground truth rotation angles.

Our pipeline includes two binary image classifiers, implemented using TensorFlow’s Keras Sequential model (TensorFlow, 2022; Table 6, in the Supporting Information), each trained using the same 80-20-10 split as the label detection model. The first, the Handwritten/Printed Classifier (HPC), distinguishes labels based on the dominant script type. The second, the Specimen Identifier Classifier (SIC), detects MfN-specific QR-code identifiers (Fig. 2, for example). After classification, SLIs are sorted by predicted class and confidence score. Classifier performance is evaluated using ground truth (GT) labels from the 10% test sets (Tables 2 and 6, in the Supporting Information), with accuracy calculated via ELIÉs label classification accuracy module.

A key challenge with the MfN_LEP_SEASIA dataset (Table 4) is the high number of images with empty labels that must be filtered out. The empty label detection module addresses this by cropping SLI images with OpenCV (Zelinsky, 2009) and analyzing the proportion of dark pixels, assuming the text is darker than the label background. This effectively removes labels without text (Table 7). Since this module uses pixel analysis rather than a trained model, a specialized testing method was applied, randomly selecting 10% of the MfN_LEP_SEASIA SLIs, yielding 2,584 unseen images representing diverse label appearances for evaluation.

**Table 7.** Performance metrics of the empty label detection module on the MfN_LEP_SEASIA test set (10% split). Runtime was measured on an Apple MacBook with an M1 CPU.

| Dataset name | Total images in test set (10% split) | Runtime (min: sec) | Average runtime per image (sec) | Accuracy (%) | Label class | Total Images Testing | Precision | Recall | F1-score |
| --- | --- | --- | --- | --- | --- | --- | --- | --- | --- |
| MfN_LEP_SEASIA | 2584 | 2:33 | 0.12 | 100 | empty | 49 | 1.00 | 1.00 | 1.00 |
|  |  |  |  |  | not_empty | 2535 | 1.00 | 1.00 | 1.00 |

OCR algorithms are optimized for horizontally aligned text, and unexpected orientations may result in decreased accuracy (Fateh et al., 2023). Hence, we implemented a rotation module on SLI in the final step of the image processing stage. A total of 1,716 images were rotated (0°, 90°, 180°, and 270°) and used to train the module with TensorFlow and Keras, using a ResNet50 base model to predict angles and rotate the image to a 0° angle (Table 3, in the Supporting Information).

### Stage 2: Text Extraction

Two different OCR platforms were used to extract textual information from images. Tesseract, an open-source OCR engine, was integrated into the pipeline using the pytesseract Python wrapper (Lee, 2023). Its performance was optimized by configuring key settings: Page Segmentation Mode 6 and Optical Engine Mode 3 (Tesseract OCR, 2024). The image preprocessing stage of the module reduces computational load and enhances image quality (Fig. 5, in the Supporting Information). Skew angle estimation, limited to -10 to 10 degrees, corrects any slant in the image, while morphological operations such as dilation and erosion remove noise that could cause unwanted results from Tesseract.

While Tesseract operates as a standalone library that requires local installation and configuration, the Google Cloud Vision API (Application Programming Interface) offers a fully managed, cloud-based solution powered by Google’s infrastructure and machine learning capabilities (Google Cloud, 2024). Using the pipeline’s Google Vision module, OCR results from the Vision API are processed not only to extract the transcribed text but also to capture bounding box information for individual words or text regions, providing valuable spatial context for the detected text.

The evaluation of the OCR systems’ performance was conducted on three datasets (Table 4): USNM_COL_CAM, MFN_LEP_SEASIA, and MCZ_ENT_Boston, using WER and CER as evaluation metrics. These metrics were computed by comparing the OCR-generated labels’ transcriptions with a manually constructed GT dataset, employing the Levenshtein distance to account for insertions, deletions, and substitutions.

To support reliable OCR performance measurement, a GT dataset was created using a clustering-based sampling method. OCR-generated transcriptions were grouped using k-medoid clustering based on Levenshtein distance, with the optimal similarity threshold selected by evaluating cluster cohesion and separation. From each multi-label cluster, the two most dissimilar labels (based on Levenshtein distance) were chosen to maximize diversity. Single-label clusters were included directly. These selected labels were then manually transcribed verbatim on the Zooniverse platform (Zooniverse, 2025), forming a representative and diverse GT dataset. This curated set was then used to compute standard OCR performance metrics such as CER and WER.

In the final step of text extraction, OCR outputs undergo post-processing to improve quality and clustering accuracy. This includes filtering out empty or implausible transcripts (such as those with extremely short average token lengths or dominated by non-alphabetic characters) based on average token length using the NLTK library in Python (Bird et al., 2009) and flagging entries like URLs. Error correction is handled through regex-based cleaning that removes non-ASCII characters, symbols, and formatting noise (e.g., pipes, degree signs, trailing punctuation). The processed transcripts are stored separately and used to extract unique tokens for the downstream clustering analysis.

### Stage 3: Text Analysis

In the final stage, the content of the corrected transcripts is analyzed in the Python-mfnb clustering package, part of the ELIE pipeline. This toolkit groups SLI text outputs with similar text content and compares the resulting clusters to assess consistency. The clustering module offers three different clustering methods: Levenshtein distance, k-medoids, and the Neighbor-Joining approach. Before grouping labels based on similarity, the transcripts are simplified by converting characters to lowercase and trimming accents.

The clustering process consists of two steps. First, seed-based clustering is performed by conducting a full-text search for each label across the entire label collection, allowing similar labels to be aggregated into clusters. Second, pairwise Levenshtein distances are calculated across labels within each cluster, and incremental partitioning is tested to improve homogeneity. Partitioning is performed using k-medoids clustering with two to 20 partitions, with the best partitioning identified by the elbow of the error sum of squares (SSE) curve. Partitioning is skipped for clusters with fewer than eight labels or non-convex SSE curves.

To assess clustering performance, four accuracy metrics were employed: the Silhouette Score, the Davies-Bouldin Index, the Calinski-Harabasz Index, and the Elbow Method. These metrics provided a comprehensive evaluation of clustering quality by measuring intra-cluster compactness, inter-cluster separation, and the overall balance of cluster structure (Rubiños et al., 2024).

To evaluate the quality of clustering, we analyzed the relationship between the OCR-generated label clusters and the GT groupings. Clusters in which all label pairs exceeded a predefined similarity threshold were provisionally marked as true positives. However, this approach can obscure over-clustering, where a single GT label is fragmented across multiple small, high-purity clusters, leading to artificially inflated accuracy scores. To address this, clustering performance was also assessed using pairwise validation, considering whether label pairs from the same actual class were correctly grouped (true positives), incorrectly split (false negatives), or merged (false positives). Clustering accuracy was evaluated across thresholds ranging from 0.3 to 0.9, using Levenshtein distance to quantify textual similarity. This iterative assessment supports the identification of an optimal threshold that balances precision and recall, minimizing both over- and under-clustering effects.

### Pipeline performance assessment

Three datasets were used to evaluate pipeline performance (Table 4): (1) MfN_LEP_SEASIA – 25,844 SLI of Pyraloidea specimens from Southeast Asia (MfN). (2) MCZ_ENT_Boston – 1,596 MLI of terrestrial insects from Boston Harbor (MCZ). (3) USNM_COL_CAM – 912 MLI of various beetle families from Central and South America (USNM).

The pipeline’s modular design enables flexible processing based on dataset characteristics and available resources. Although all modules can be run universally, selective use reduces processing time, important for users with limited computing power. All tests were run on an Apple M1 CPU.

1. MfN_LEP_SEASIA used seven modules (excluding Label Detection due to SLI): Empty Label Detection, SIC, HPC, Label Rotation, Google Vision OCR, Label Post-processing, and Clustering, with a runtime of 2 hours and 44 minutes.
2. MCZ_ENT_Boston used five modules: Label Detection, HPC, Google Vision OCR, Label Post-processing, and Clustering. Modules for empty labels, specimen IDs, and rotation were skipped due to dataset characteristics. It had a runtime of 51 minutes and 10 seconds.
3. USNM_COL_CAM ran all eight modules, with a runtime of 47 minutes and 21 seconds.

To visualize clustering results, a pre-trained Gensim Word2Vec model (Rehurek & Sojka, 2011) (Fig. 6) generated 100-dimensional embeddings for each label. These were reduced to 2D using t-SNE (Maaten & Hinton, 2008) and plotted with Plotly Express (Plotly, 2015). Each dot represents a label, colored by its assigned cluster, enabling comparison of clustering outputs and semantic similarity.

**Figure 6.**
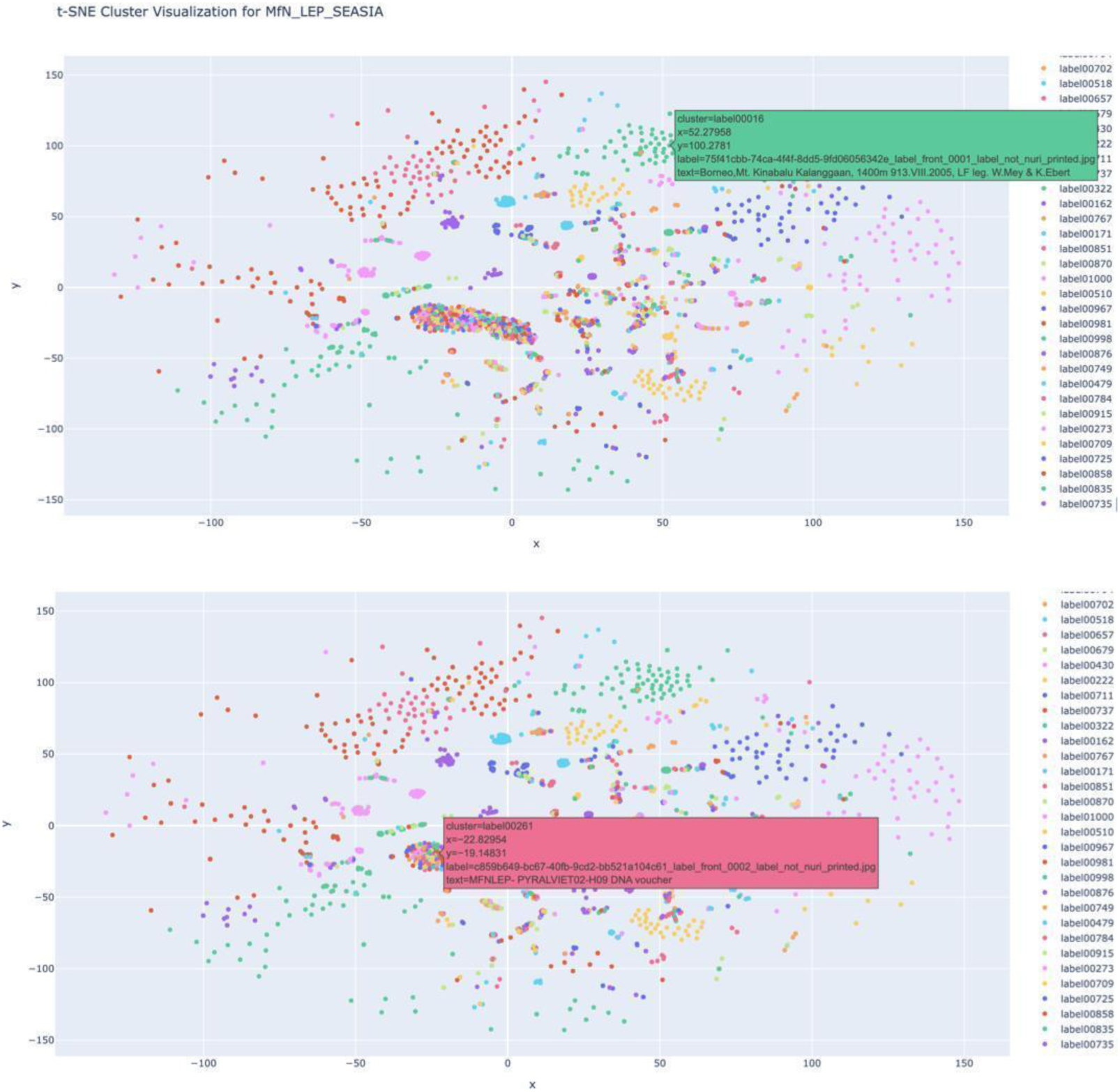
Two-dimensional t-SNE visualization of multi-label clusters in the MfN_LEP_SEASIA dataset. Each point represents a single-label image (SLI) transcript, converted into a 100-dimensional vector using a pre-trained Gensim Word2Vec model. These vectors were averaged across tokens per label and reduced to two dimensions via t-SNE (perplexity = 30, n_iter = 1,000). Colors indicate the cluster assignments based on Levenshtein-distance clustering at a 0.9 similarity threshold. Tight spatial groupings reflect labels with highly similar content, while dispersed points suggest either semantically distinct labels or OCR artifacts. The figure was rendered with Plotly Express for a visual comparison of embedding and clustering results.

## Results

### Label Detection

The label detection module processed 278 images from the test set in 4 minutes and 46 seconds (1.03 sec./image, Table 5, in the Supporting Information), achieving a 94% accuracy rate. Each output includes the predicted class ("label"), bounding box coordinates, and a confidence score ranging from 0 to 1. A customizable score threshold of 0.8 is applied to filter out low-confidence predictions, reducing false positives and false negatives (Fig. 4.1). The box plot shows that most IOU scores exceed 0.8, indicating strong alignment between predicted and annotated bounding boxes.

### Empty Label Detection

The empty label detection module was applied to 10% of the MfN_LEP_SEASIA dataset: 2,584 SLI images. The performance of the module was evaluated using an accuracy test to assess text density predictions, and runtime calculations were recorded (Table 7). The model achieved perfect precision (1.00) for detecting non-empty labels, and the high recall score indicates that the module successfully detects all non-empty labels without false negatives.

### Label Classification

The SIC achieves maximum performance with a score of 1.00 across all metrics for both "identifier" and "not_identifier" categories (Table 8 and Fig. 4.3). HPC demonstrates substantial precision and recall metrics, with handwritten text demonstrating a better recall at 0.99 compared to 0.98 for printed text (Fig. 4.2).

**Table 8.** Performance metrics of the label classification module for two binary classification models. Runtimes were measured on an Apple MacBook with an M1 CPU.

| Model Name | Classes | Runtime (sec) | Avg. runtime per image (sec) | Accuracy (%) | Precision | Recall | F1-score |
| --- | --- | --- | --- | --- | --- | --- | --- |
| Specimen Identifier Classifier | identifier | 0.34 | 0.85 | 100 | 1.00 | 1.00 | 1.00 |
|  | not_identifier |  |  |  | 1.00 | 1.00 | 1.00 |
| Handwritten Printed Classifier | handwritten | 0.55 | 0.86 | 99 | 0.98 | 0.99 | 0.99 |
|  | printed |  |  |  | 0.99 | 0.98 | 0.98 |

### Label Rotation

The label rotation module handled the test set, comprising 156 images (Table 3, in the Supporting Information), completing the task in 7.4 seconds (0.05 sec./per image). After comparing the predictions to the GT, the model achieved an accuracy of 95% (Table 9). The analysis of the confusion matrix reveals specific misclassifications across different rotation angles (Fig. 4.4). Notably, all four incorrectly assigned images were misclassifications as ±180° from the proper angle.

**Table 9.** Performance metrics of the label rotation classification module at four angle orientations. Runtimes were measured on an Apple MacBook with an M1 CPU.

| Angles | Runtime (sec) | Avg. runtime per image (sec) | Accuracy (%) | Precision | Recall | F1-score |
| --- | --- | --- | --- | --- | --- | --- |
| 0° | 7.28 | 0.05 | 95 | 0.95 | 0.95 | 0.95 |
| 90° |  |  |  | 0.95 | 0.95 | 0.95 |
| 180° |  |  |  | 0.95 | 0.95 | 0.95 |
| 270° |  |  |  | 0.95 | 0.95 | 0.95 |

### OCR Evaluation

While a comprehensive OCR software evaluation was beyond this study’s scope, the Google Cloud Vision API significantly outperformed Tesseract across all datasets. Google Vision demonstrated strong performance, particularly on the MCZ_ENT_Boston dataset, where it achieved a median WER and CER of 0.00. In contrast, Tesseract’s performance was weaker and more variable, struggling most notably with the MCZ_ENT_Boston dataset, where its mean CER exceeded 1.2. The detailed error distributions for both OCR systems across all three test datasets are presented in Figures 7 and 8.

**Figure 7.**
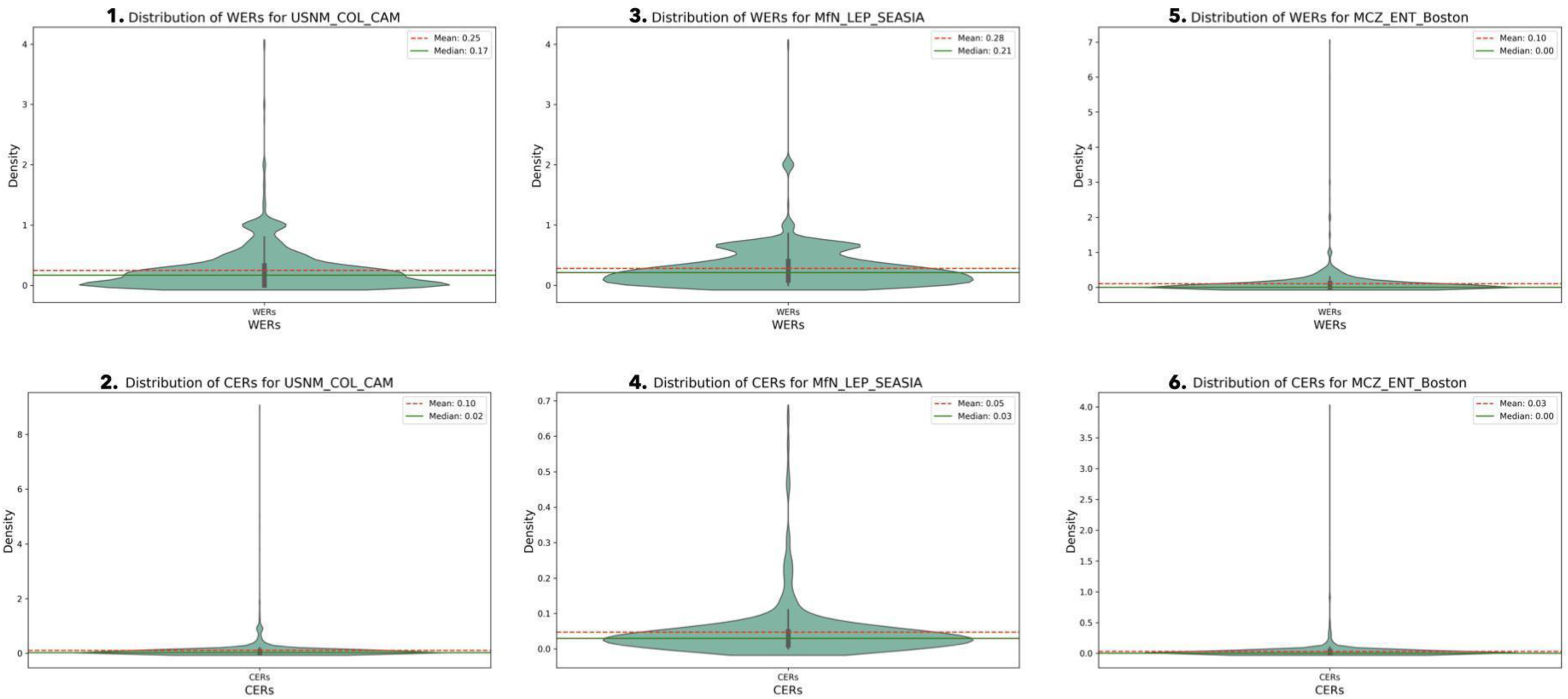
Distribution of OCR error rates produced by the Google Vision API across the three testing datasets. Violin plots display the distributions of Character Error Rate (CER) and Word Error Rate (WER) per label, evaluated against manually transcribed ground truth. Lower values indicate better OCR performance. Each violin reflects the density of error rates for individual labels; horizontal lines represent the median and the mean. CER captures the percentage of incorrect characters, while WER measures word-level errors, including insertions, deletions, and substitutions.

**Figure 8.**
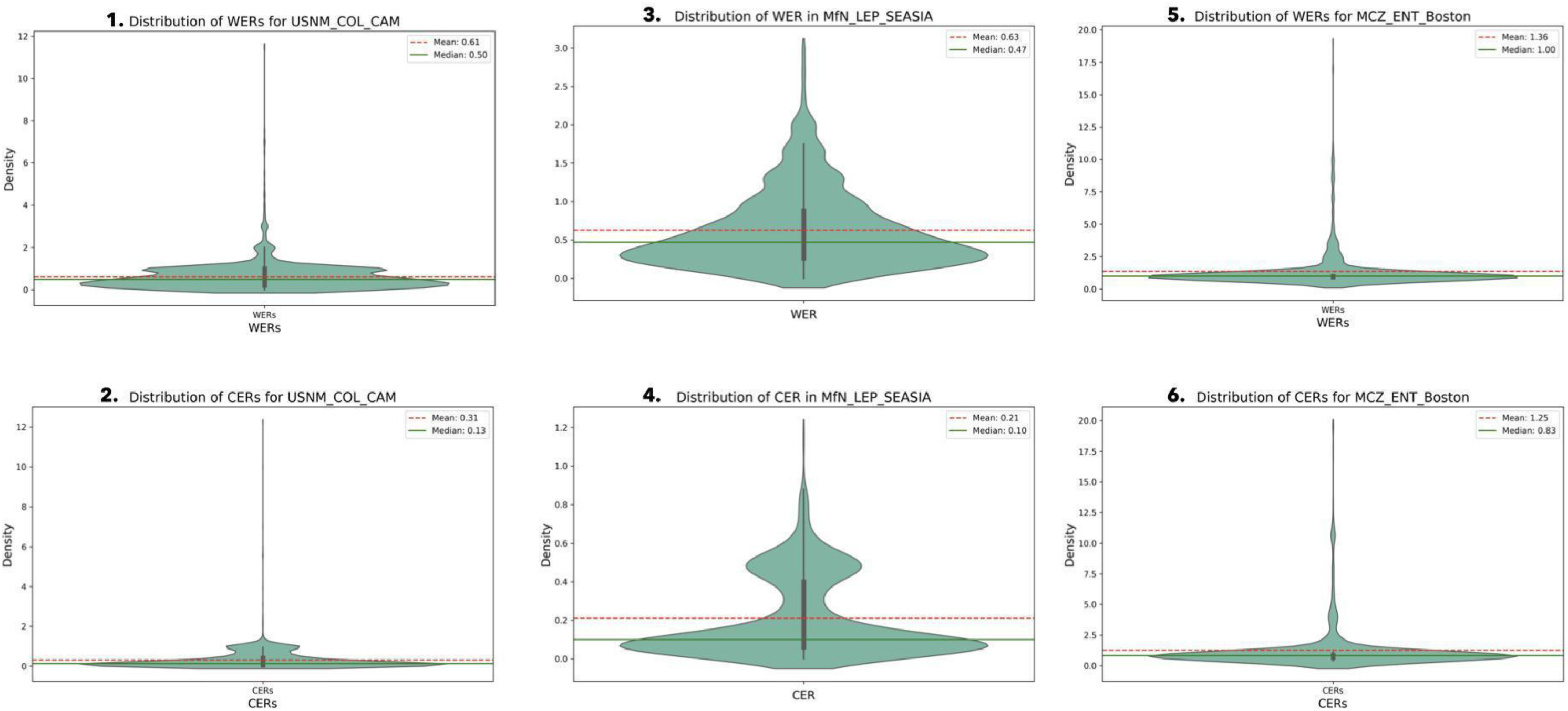
Distribution of OCR error rates produced by Tesseract across the three testing datasets. Violin plots display the distributions of Character Error Rate (CER) and Word Error Rate (WER) per label, evaluated against manually transcribed ground truth. Lower values indicate better OCR performance. Each violin reflects the density of error rates for individual labels; horizontal lines represent the median and the mean. CER captures the percentage of incorrect characters, while WER measures word-level errors, including insertions, deletions, and substitutions.

### Label Clustering

Our clustering analysis revealed distinct performance characteristics across the datasets. For MfN_LEP_SEASIA, clustering performance was strong, yielding compact and well-separated groups (Fig. 9.1). This was evidenced by high Silhouette Scores and a steadily increasing Calinski-Harabasz Index, which peaked at 0.9 similarity threshold (Fig. 10.1). At this optimal threshold, a high proportion of labels were successfully grouped into meaningful clusters (Table 10.1, in the Supporting Information).

**Figure 9.**
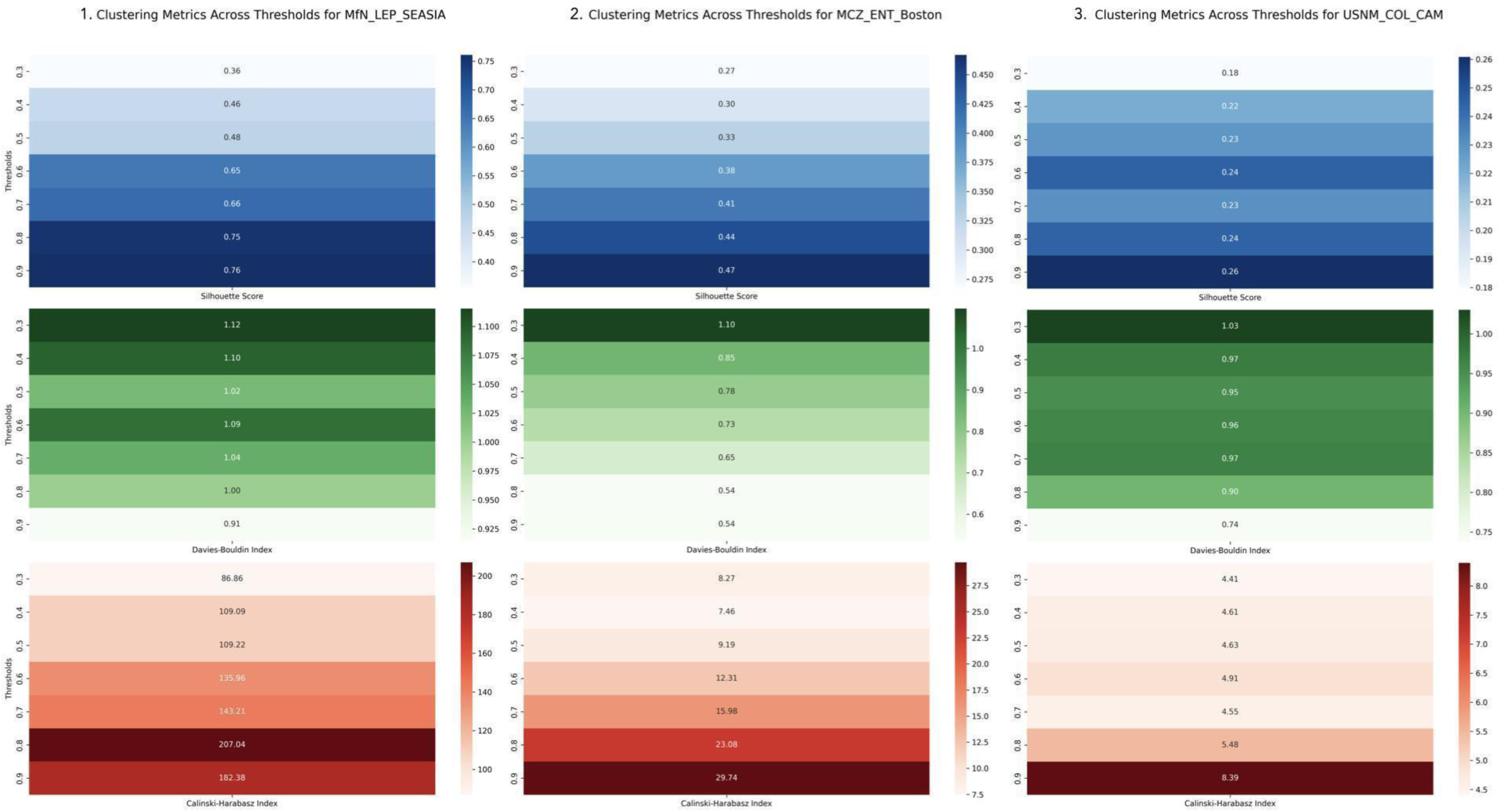
Clustering Accuracy Metrics Across Similarity Thresholds. Performance of the clustering module for the three test datasets across similarity thresholds from 0.3 to 0.9. Each line represents one clustering evaluation metric: Silhouette Score (cohesion and separation), Davies-Bouldin Index (intra/inter-cluster similarity), and Calinski-Harabasz Index (cluster dispersion). Higher Silhouette and Calinski-Harabasz scores, and lower Davies-Bouldin scores, indicate better clustering.

**Figure 10.**
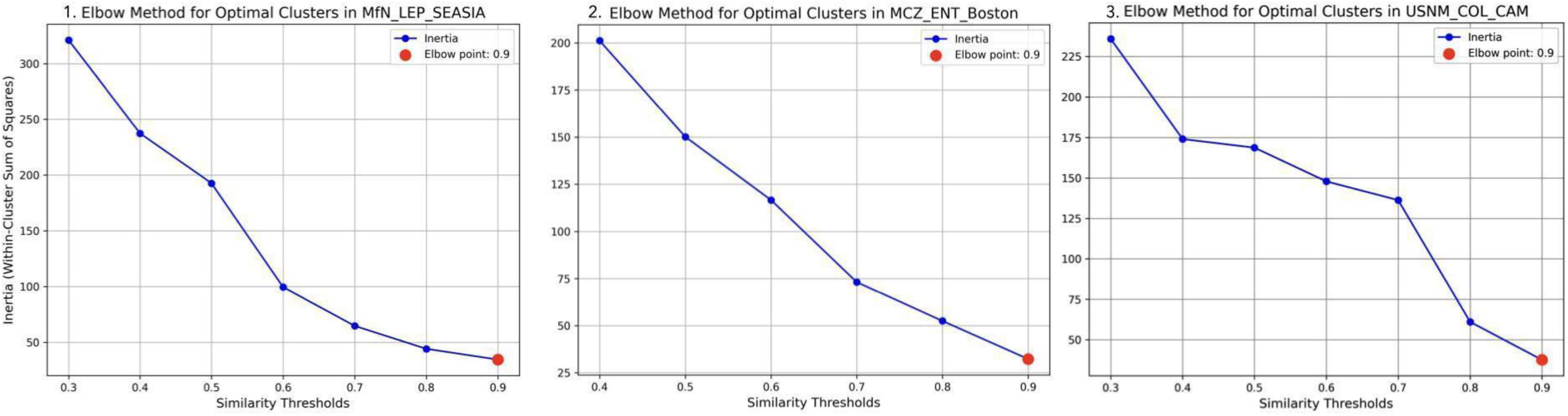
Clustering Optimization via Elbow Method Across Datasets. Elbow method plots were used to determine the optimal number of clusters at each threshold for the three test datasets: (1) MfN_LEP_SEASIA, (2) MCZ_ENT_Boston, and (3) USNM_COL_CAM. The number of clusters (x-axis) is plotted against the sum of squared errors (SSE; y-axis) for k-medoids clustering, showing diminishing returns beyond an inflection point (the “elbow”). The elbow point (typically around threshold = 0.9) indicates the best balance between cluster compactness and separation.

Similarly, USNM_COL_CAM showed improved clustering with higher thresholds, with the elbow method confirming 0.9 as optimal (Figs. 9.3 and 10.3). While its clustering metrics were slightly lower than MfN_LEP_SEASIA, indicating less uniform clusters, the pipeline still effectively grouped a significant number of labels (Table 10.2, in the Supporting Information).

In contrast, MCZ_ENT_Boston exhibited weaker clustering performance (Fig. 9.2). Its lower Calinski-Harabasz Index and moderate Silhouette Score reflect minimal variation between labels, often differing only by minor details like GPS coordinates or date. This resulted in a higher number of small or overlapping clusters, particularly at lower thresholds, suggesting limited distinctiveness in the data (Table 10.3, in the Supporting Information). Despite this, 0.9 remained the optimal threshold (Fig. 10.2).

### Clustering Validation

At the optimal similarity threshold of 0.9, clustering accuracy varied across datasets (Fig. 11). The USNM_COL_CAM dataset achieved the highest accuracy at 98%, with only 27 false-positive clusters and minimal over-splitting (0.8% false negatives). The MfN_LEP_SEASIA dataset yielded a clustering accuracy of 79.1%, where 91 clusters were identified as false positives and 24% of clusters were affected by over-splitting. Performance on the MCZ_ENT_Boston dataset was similar, with an accuracy of 70.6%, 67 false-positive clusters, and only 1.6% of labels impacted by over-splitting. Box plots of the internal Levenshtein distances for each dataset’s clusters are shown in Figure 12.

**Figure 11.**
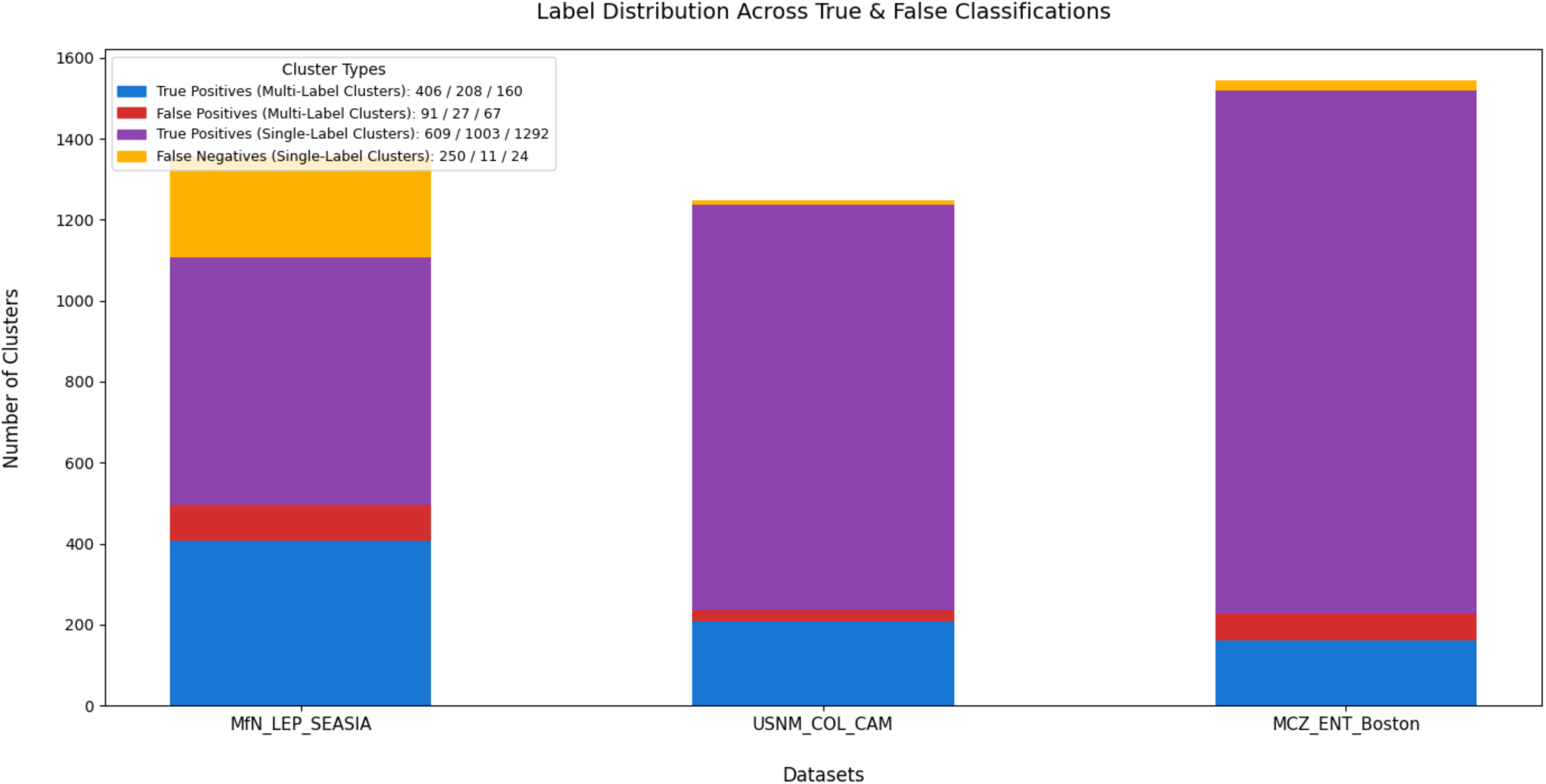
Clustering validation results across datasets and similarity thresholds. Bars represent the number of true positives, false positives, and false negatives resulting from the pairwise validation of clustering outcomes against ground truth label groupings. Results are shown separately for the three datasets: MfN_LEP_SEASIA, USNM_COL_CAM, and MCZ_ENT_Boston. Clustering was performed at a similarity threshold of 0.9 using Levenshtein distance. While high clustering accuracy was achieved in MfN_LEP_SEASIA and USNM_COL_CAM, over-splitting (false negatives) and occasional label misgroupings (false positives) were more frequent in MCZ_ENT_Boston due to higher text similarity across labels.

**Figure 12.**
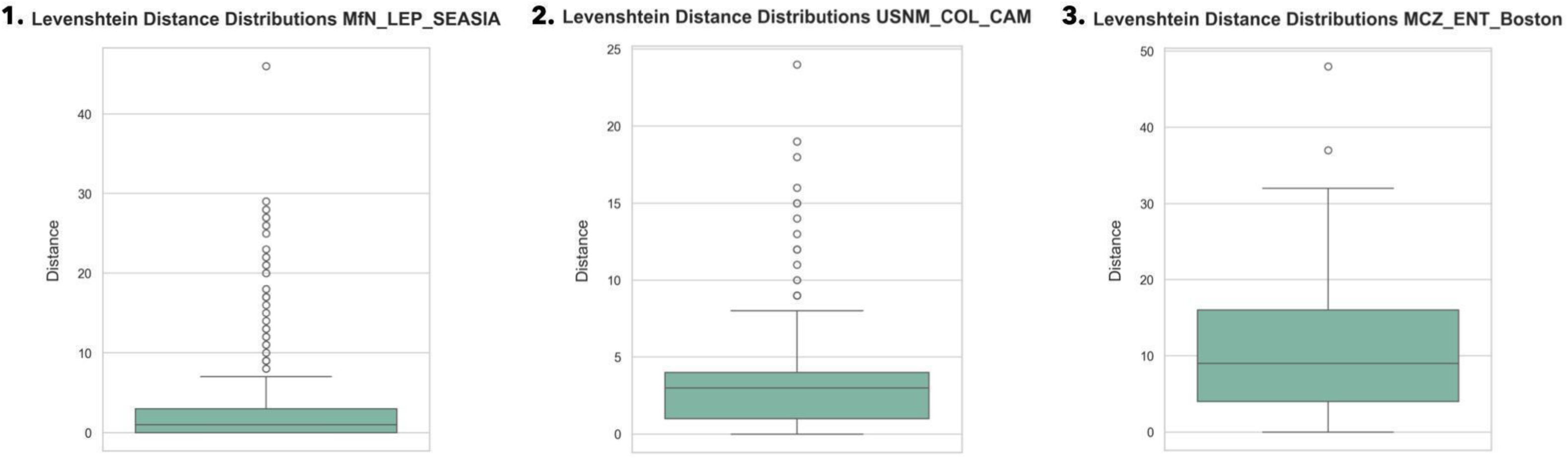
Distribution of the maximum Levenshtein distances between label pairs within each multi-label cluster at a similarity threshold of 0.9. Each subplot represents one dataset: (1) MfN_LEP_SEASIA, (2) USNM_COL_CAM, and (3) MCZ_ENT_Boston. Lower distances indicate a higher internal consistency of clusters. Outliers suggest false positives (non-identical labels grouped). Histograms are truncated at the 95th percentile to improve visualization.

## Discussion

### Streamlining Label Information Extraction

The presented pipeline efficiently handles images from large-scale digitization projects by rapidly processing redundant labels and reserving challenging ones for detailed manual review. The mitigation of performance issues in HTR models due to handwriting variability and limited training data (Ströbel et al., 2022) was the rationale for separating handwritten and printed text in our pipeline, allowing each to be processed with specialized CNN models. While historical labels often contain handwritten text, more recent ones almost exclusively contain printed text. By accounting for 94% of the total labels in the MfN_LEP_SEASIA dataset, 72% in USNM_COL_CAM, and 84% in MCZ_ENT_Boston, printed labels represent a major part of the datasets investigated here. However, this reflects a bias stemming from the relatively recent origin of the specimens investigated, whereas other collections contain specimens primarily dating from the late 19th to early 20th centuries (e.g., Garner et al., 2024).

Manual transcription and inference of metadata such as geolocalisation remain time-consuming, averaging 1.46 minutes for handwritten and 4.80 minutes for printed labels (Walton et al., 2020a). Because entomological labels frequently contain repeated information, for instance, collection data or taxonomic identification, capturing each unique label once and linking its duplicates is the most efficient approach.

Lower clustering thresholds can furthermore help to group semantically similar labels. For example, clustering the labels of the MfN_LEP_SEASIA dataset at a threshold of 0.3 sorted the labels in groups of identical categories, such as collecting event, specimen identifier, taxon ID, collection, DNA, type, preparation, enabling a presorting of similar labels for a more efficient data validation.

### Image Quality and OCR Performance

Image and label quality influence OCR accuracy in our pipeline. High-resolution images with strong contrast yield better results, while low-resolution scans, poor lighting, and physical label imperfections often reduce performance. For example, even high-resolution labels like "GU SP 409 ♀" (Fig. 13.1, in the Supporting Information) produced high error rates (WER/CER: 0.67) due to uneven inking and spacing. Similarly, labels with poor contrast (e.g., dark backgrounds, complex fonts) or irregular formatting, like hand-cut edges or blurred text (Figs. 13.2, 13.3, 13.4, in the Supporting Information), led to substantial recognition errors.

These examples underscore the challenge of extracting text from legacy labels not designed for machine readability. While standardized imaging can mitigate some issues, it cannot fully resolve problems inherent to deteriorated or non-standard labels.

Among OCR tools, Google Cloud Vision API consistently outperformed Tesseract across all datasets. While Tesseract achieved acceptable character accuracy, it struggled with word segmentation and complex layouts. Google Vision showed greater robustness in handling variability in text and formatting, particularly in datasets like MfN_LEP_SEASIA and MCZ_ENT_Boston.

Integrating OCR services such as Microsoft Azure or Amazon Textract could offer benefits like multi-language support, layout-aware extraction, and improved handling of mixed content (e.g., Latin taxonomy and vernacular terms).

### Modular Architecture, Scalability, and Cross-Institutional Applicability

The modular architecture of our pipeline allows flexible deployment based on dataset complexity. Clean datasets (i.e., exempt from empty labels) can skip steps like Empty Label Detection or Label Rotation, while more complex workflows benefit from full execution, including SIC and HPC modules. Parallel processing enables concurrent execution across computational nodes, making our approach scalable for both small collections and large institutional repositories.

Designed for cross-institutional use, our pipeline has been trained and tested on diverse datasets, from community platforms like AntWeb to institutional efforts such as *digitize!* (MfN, 2025), and collections from USNM as well as MCZ. Its compatibility with varied imaging protocols, including legacy data not originally designed for AI integration (Dodge & Karam, 2016), makes it adaptable to heterogeneous digitization contexts. Comprehensive guidelines and training scripts are provided, allowing the models to be retrained on specific datasets. The architecture also supports extensibility, enabling integration of future modules such as Named Entity Recognition (NER), Large Language Models (LLMs), or HTR.

### Refining the Pipeline: Recommendations and Future Directions

While our pipeline provides a strong baseline for extracting insect label data, several improvements can be made. First, combining preprocessing techniques, following OCR-D project insights (OCR-D Coordination Project, 2023), could boost OCR accuracy. Incorporating advanced object detection models such as Detectron2 (Wu et al., 2019) could improve label segmentation, especially for complex layouts. To better handle handwritten data, which remains challenging for OCR, developing a dedicated HTR module using tools like Transkribus would extend support for historical labels (Hardisty & Roberts, 2013). Developing a user interface would make our approach more accessible for non-technical users. Finally, developing downstream tools for language identification and NER will enable automatic classification of key label elements, such as specimen IDs, collection sites, and taxonomic names.

### Benefits of collection-based research

By greatly improving digitization workflows, our approach significantly enhances the scientific value of entomological collections for ecological and evolutionary research. It enables the reconstruction of long-term trends in species distributions, phenology, and community composition (e.g., Bartomeus et al., 2018; Habel et al., 2016; Nufio et al., 2025) and the discovery and description of new species (e.g., Léger et al., 2024). Other insights can be gained on the taxonomical width and depth as well as the historical development of specific collections (Garner et al., 2024). Standardized, high-quality metadata for specimen-based datasets underpin analyses of range shifts, phenological change, and trait-based responses to environmental pressures (Kharouba et al. 2018). Through the semi-automated, large-scale extraction and structuring of metadata, our pipeline facilitates the transformation of insect collections into accessible, research-ready datasets, thereby enhancing the utility of museum archives for biodiversity research.

## Conclusion

ELIE (Entomological Label Information Extraction) is a modular, semi-automated pipeline for extracting and processing insect specimen label data. Using CNNs, it detects and crops labels, classifies text types, and applies tailored OCR, achieving high accuracy for printed labels. Automated extraction and clustering can reduce manual transcription by up to 90%, significantly accelerating the digitization process. Our pipeline’s flexible architecture supports future additions: HTR, NER, and LLMs. Though built for entomology, its design is adaptable to fields such as archaeology or manuscript studies, helping modernize collection workflows and expand access to cultural and scientific data.

## Acknowledgments

We are grateful to the volunteers from the Transcription Workshop team at the MfN for their invaluable work in transcribing the specimen labels used in this project. We gratefully acknowledge the financial support provided by the Internal Innovation Fund of the MfN, which enabled Joël Tuberosa to develop the clustering module. We are especially thankful to Bonnie Blaimer (MfN), Brian Fisher (AntWeb), and Crystal Maier (MCZ) for generously sharing their image datasets. Our thanks also go to Joachim Snellings (MfN), Conor Fahy (MfN), and Majid Vafadar (MfN) for their technical contributions to the development of the pipeline, and to Frederik Berger (MfN) for his continued logistical support throughout the project.

## Data Availability

● Training, validation, and testing datasets (including entomological label images) are archived on Zenodo under the project “ELIE - Entomological Label Information Extraction”: https://doi.org/10.7479/khac-x956 (Belot, 2025).
● The complete source code for the ELIE pipeline, including pre-trained models and workflow scripts, is available under the GNU General Public License v3.0 and hosted on GitHub: https://github.com/MargotBelot/entomological-label-information-extraction.git.
● The python-mfnb package, which provides clustering algorithms and associated reference data, is also released under the GNU General Public License v3.0 and can be found on GitHub: https://github.com/MargotBelot/mfnb_clustering.git.
● User documentation, including installation instructions, module descriptions, and example workflows, is accessible via ReadTheDocs: https://python-label-processing.readthedocs.io/en/latest/index.html.

## Inclusion and Diversity Statement

This project was conducted by a gender-balanced team of researchers based at the Museum für Naturkunde in Berlin. The team represents a range of career stages, from students and early-career researchers to senior scientists, and is committed to fostering an inclusive research environment. The project promoted open science by using open-source software, publishing openly accessible data, and engaging the public through citizen science contributions on the Zooniverse platform. These efforts support broader participation and equitable access to biodiversity knowledge.

## Author Contributions

Margot Belot developed the pipeline, implemented core modules, performed all testing and performance evaluations, and wrote the manuscript. Joël Tuberosa developed the clustering module, supported evaluation, and contributed to writing the clustering section of the manuscript. Leonardo Preuss co-developed all modules. Olga Svezhentseva contributed to the development of the OCR and post-processing modules. Magdalena Claessen assisted with dataset curation and image preprocessing. Christian Bölling and Franziska Schuster contributed to the initial setup and technical infrastructure of the project. Théo Léger conceived and led the project and co-wrote the manuscript. All authors reviewed and approved the final version of the manuscript.

## References

Ahrens, D., Haas, A., Pacheco, T.L. & Grobe, P. (2025). “Extracting specimen label data rapidly with a smartphone - a great help for simple digitization in taxonomy and collection management.” ZooKeys, 1233, pp.15–30. 10.3897/zookeys.1233.140726.

California Academy of Sciences. (n.d.). AntWeb. [online] Available at: https://www.antweb.org/ [Accessed 4 Jul. 2025].

Arce, A.N., Cantwell-Jones, A., Tansley, M., Barnes, I., Brace, S., Mullin, V.E., Notton, D. et al. (2022). “Signatures of increasing environmental stress in bumblebee wings over the past century: Insights from museum specimens.” Journal of Animal Ecology, 92(2), pp.297–309. 10.1111/1365-2656.13788.

Bartomeus, I., Stavert, J.R., Ward, D. & Aguado, O. (2018). “Historical collections as a tool for assessing the global pollination crisis.” Philosophical Transactions of the Royal Society B, 374(1763), p.20170389. 10.1098/rstb.2017.0389.

Belot, M. (2025). ELIE - Entomological Label Information Extraction [Data set]. Museum für Naturkunde (MfN) - Leibniz-Institut für Evolutions- und Biodiversitätsforschung. 10.7479/khac-x956.

Bi, A. (2022). “Detecto (1.2.2).” Detecto: Computer Vision and Object Detection Models. [online] Available at: https://pypi.org/project/detecto/. [Accessed 05 Jun. 2025].

USDA Agricultural Research Service. (n.d.). Big-Bee Project. [online] Available at: http://big-bee.net/ [Accessed 4 Jul. 2025].

Bird, S., Klein, E. & Loper, E. (2009). Natural language processing with Python. O’Reilly Media.

Blagoderov, V., Kitching, I., Livermore, L., Simonsen, T. & Smith, V. (2012). “No specimen left behind: Industrial scale digitization of natural history collections.” ZooKeys, 209, pp.133–146. 10.3897/zookeys.209.3178.

Brooks, S.J., Self, A., Toloni, F. & Sparks, T. (2014). “Natural history museum collections provide information on phenological change in British butterflies since the late-nineteenth century.” International Journal of Biometeorology, 58(8), pp.1749–1758. 10.1007/s00484-013-0780-6.

Cridland, J.M., Ramirez, S.R., Dean, C.A., Sciligo, A. & Tsutsui, N.D. (2018). “Genome sequencing of museum specimens reveals rapid changes in the genetic composition of honey bees in California.” Genome Biology and Evolution, 10(2), pp.458–472. 10.1093/gbe/evy007.

Plotly. (2015). “Data Apps for Production.” [online] Available at: https://plot.ly/. [Accessed 05 Jun. 2025].

De Araújo Romeiro, L., Borges, R.C., Da Silva, E.F., Guimarães, J.T.F. & Giannini, T.C. (2023). “Assessing entomological collection data to build pollen interaction networks in the tropical Amazon forest.” Arthropod-Plant Interactions, 17(3), pp.313–325. 10.1007/s11829-023-09968-7.

Garner, B.H., Creedy, T.J., Allan, E.L., Crowther, R., Devenish, E., Kokkini, P., Livermore, L., Lohonya, K., Lowndes, N., Wing, P., and Vogler, A.P. (2024). The taxonomic composition and chronology of a museum collection of Coleoptera were revealed through large-scale digitisation. Front. Ecol. Evol., 17 July 2024, Sec. Phylogenetics, Phylogenomics, and Systematics Volume 12. 10.3389/fevo.2024.1305931

Museum für Naturkunde Berlin (MfN). (2025). Digitize Project. [online] Available at: https://www.museumfuernaturkunde.berlin/en/museum/exhibitions/digitize. [online] [Accessed 4 Jul. 2025].

Dodge, S. & Karam, L. (2016). “Understanding how image quality affects deep neural networks.” 8th International Conference on Quality of Multimedia Experience (QoMEX), pp.1–6. 10.1109/qomex.2016.7498955.

Fateh, A., Fateh, M. & Abolghasemi, V. (2023). “Enhancing optical character recognition: Efficient techniques for document layout analysis and text line detection.” Engineering Reports, 6(9). 10.1002/eng2.12832.

Fontaine, B., Perrard, A. & Bouchet, P. (2012). “21 years of shelf life between discovery and description of new species.” Current Biology, 22(22), pp.R943–R944. 10.1016/j.cub.2012.10.029.

Freedman, M.G., Dingle, H., Strauss, S.Y. & Ramírez, S.R. (2020). “Two centuries of monarch butterfly collections reveal contrasting effects of range expansion and migration loss on wing traits.” Proceedings of the National Academy of Sciences, 117(46), pp.28887–28893. 10.1073/pnas.2001283117.

Gotelli, N.J., Booher, D.B., Urban, M.C., Ulrich, W., Suarez, A.V., Skelly, D.K., Russell, D.J. et al. (2021). “Estimating species relative abundances from museum records.” Methods in Ecology and Evolution, 14(2), pp.431–443. 10.1111/2041-210x.13705.

Habel, J. C., Segerer, A., Ulrich, W., Torchyk, O., Weisser, W. W., & Schmitt, T. (2016). Butterfly community shifts over two centuries. Conservation Biology, 30(4), 754–762. 10.1111/cobi.12656

Halsch, C.A., Shapiro, A.M., Fordyce, J.A., Nice, C.C., Thorne, J.H., Waetjen, D.P. & Forister, M.L. (2021). “Insects and recent climate change.” Proceedings of the National Academy of Sciences, 118(2). 10.1073/pnas.2002543117.

Hardisty, A. & Roberts, D. (2013). “A decadal view of biodiversity informatics: Challenges and priorities.” BMC Ecology, 13(1), p.16. 10.1186/1472-6785-13-16.

Ingle, R.R., Fujii, Y., Deselaers, T., Baccash, J. & Popat, A.C. (2019). “A scalable handwritten text recognition system.” 2019 International Conference on Document Analysis and Recognition (ICDAR), pp.17–24. 10.1109/icdar.2019.00013.

Smithsonian Institution. (2025). “National Collections.” Available at: https://www.si.edu/dashboard/national-collections#collections-digitization. [online] [Accessed 28 Jan. 2025].

Kharouba, H.M., Lewthwaite, J.M.M., Guralnick, R., Kerr, J.T. & Vellend, M. (2018). “Using insect natural history collections to study global change impacts: challenges and opportunities.” Philosophical Transactions of the Royal Society B, 374, p.20170405. 10.1098/rstb.2017.0405.

Kirchhoff, A., Bügel, U., Santamaria, E., Reimeier, F., Röpert, D., Tebbje, A., Güntsch, A. et al. (2018). “Toward a service-based workflow for automated information extraction from herbarium specimens.” Database, 2018. 10.1093/database/bay103.

PyPI. (2021). “LabelImg.” Available at: https://pypi.org/project/labelImg/. [online] [Accessed 05 Jun. 2025].

Lee, M. (2023). “GitHub - Madmaze/Pytesseract: A Python Wrapper for Google Tesseract.” Available at: https://github.com/madmaze/pytesseract. [online] [Accessed 05 Jun. 2025].

Léger, T. (2024). Half of the Diversity Undescribed: Integrative Taxonomy Reveals 32 New Species and a High Cryptic Diversity in the Scopariinae and Crambinae of the Philippines (Lepidoptera: Crambidae). Bulletin of the Society of Systematic Biologists, 3(2). 10.18061/bssb.v3i2.9527.

Lepidoptera of North America Network. (n.d.). LepNet. [online] Available at: http://www.lep-net.org/ [Accessed 4 Jul. 2025].

Maaten, L. & Hinton, G.E. (2008). “Visualizing data using t-SNE.” Available at: https://api.semanticscholar.org/CorpusID:5855042. [online] [Accessed 05 Jun. 2025].

Natural History Museum London (NHM). (2025). Data Portal. [online] Available at: https://data.nhm.ac.uk/. [online] [Accessed 4 Jul. 2025].

Nufio, C.R., Sheffer, M.M., Smith, J.M., Troutman, M.T., Bawa, S.J., Taylor, E.D., et al. (2025). “Insect size responses to climate change vary across elevations according to seasonal timing.” PLoS Biol 23(1): e3002805. 10.1371/journal.pbio.3002805

OCR-D. (2023). “OCR-D Coordination Project.” Available at: https://ocr-d.de/en. [online] [Accessed 05 Jun. 2025].

Owen, D., Groom, Q., Hardisty, A., Leegwater, T., Livermore, L., Van Walsum, M., Wijkamp, N. & Spasić, I. (2020). “Towards a scientific workflow featuring natural language processing for the digitization of natural history collections.” Research Ideas and Outcomes, 6. 10.3897/rio.6.e58030.

Pérez-Lachaud, G. & Lachaud, J.-P. (2017). “Hidden biodiversity in entomological collections: The overlooked co-occurrence of dipteran and hymenopteran ant parasitoids in stored biological material.” PLoS ONE, 12(9), e0184614. 10.1371/journal.pone.0184614.

Platts, P.J., Mason, S.C., Palmer, G., Hill, J.K., Oliver, T.H., Powney, G.D., Fox, R. & Thomas, C.D. (2019). “Habitat availability explains variation in climate-driven range shifts across multiple taxonomic groups.” Scientific Reports, 9(1). 10.1038/s41598-019-51582-2.

Ponder, W.F., Carter, G.A., Flemons, P. & Chapman, R.R. (2001). “Evaluation of museum collection data for use in biodiversity assessment.” Conservation Biology, 15(3), pp.648–657. 10.1046/j.1523-1739.2001.015003648.x.

Poske, C. (2024). “Digital repatriation of cultural heritage.” In: The Palgrave Encyclopedia of Cultural Heritage and Conflict. Cham: Springer Nature Switzerland, pp.1–5. 10.1007/978-3-030-61493-5_148-1.

Price, B.W., Dupont, S., Allan, E.L., Blagoderov, V., Butcher, A.J., Durrant, J., Holtzhausen, P. et al. (2018). “ALICE: Angled label image capture and extraction for high throughput insect specimen digitization.” OSF Preprints, 5 Nov. 10.31219/osf.io/s2p73.

Ptucha, R., Petroski Such, F., Pillai, S., Brockler, F., Singh, V. & Hutkowski, P. (2018). “Intelligent character recognition using fully convolutional neural networks.” Pattern Recognition, 88, pp.604–613. 10.1016/j.patcog.2018.12.017.

Rakosy, D., Ashman, T.-L., Zoller, L., Stanley, A. & Knight, T.M. (2022). "Integration of historical collections can shed light on patterns of change in plant-pollinator interactions and pollination service." Functional Ecology, 37(2), pp.218–233. 10.1111/1365-2435.14211.

Raschka, S. & Mirjalili, V. (2019). Python machine learning: Machine learning and deep learning with Python, scikit-learn, and TensorFlow 2. Birmingham: Packt Publishing.

Raven, P.H. & Wagner, D.L. (2021). “Agricultural intensification and climate change are rapidly decreasing insect biodiversity.” Proceedings of the National Academy of Sciences, 118(2). 10.1073/pnas.2002548117.

Rehurek, R. & Sojka, P. (2011). "Gensim–Python framework for vector space modeling." NLP Centre, Faculty of Informatics, Masaryk University, 3(2).

Rubiños, M., Díaz-Longueira, A., Timiraos, M., Michelena, Á., García-Ordás, M.T. & Alaiz-Moretón, H. (2024). “A comparative analysis of algorithms and metrics to perform clustering.” In: Lecture Notes in Networks and Systems, pp.63–72. 10.1007/978-3-031-73910-1_7.

Short, A.E.Z., Dikow, T. & Moreau, C.S. (2018). “Entomological collections in the age of big data.” Annual Review of Entomology. 10.1146/annurev-ento-031616-035536.

Stork, N.E. (2017). “How many species of insects and other terrestrial arthropods are there on Earth?” Annual Review of Entomology, 63(1), pp.31–45. 10.1146/annurev-ento-020117-043348.

Ströbel, P.B., Clematide, S., Volk, M., Schwitter, R., Hodel, T. & Schoch, D. (2022). “Evaluation of HTR models without ground truth material.” arXiv. 10.48550/arxiv.2201.06170.

Takano, A., Cole, T.C.H. & Konagai, H. (2024). “A novel automated label data extraction and database generation system from herbarium specimen images using OCR and NER.” Scientific Reports, 14(1). 10.1038/s41598-023-50179-0.

TensorFlow. (2022). “TensorFlow: A machine learning library for Python.” Available at: https://www.tensorflow.org/.

Tesseract OCR. (2024). “Tesseract user manual.” GitHub Documentation (Version 5.3.4). Available at: https://tesseract-ocr.github.io/tessdoc/.

Transkribus. (2025). Available at: https://www.transkribus.org/. [online] [Accessed 17 Apr. 2025].

Google Cloud. (2024). “Vision AI: Image & Visual AI Tools.” Available at: https://cloud.google.com/vision. [online] [Accessed 05 Jun. 2025].

Walton, S., Livermore, L., Bánki, O., Cubey, R., Drinkwater, R., Englund, M., Goble, C. et al. (2020b). “Landscape analysis for the Specimen Data Refinery.” Research Ideas and Outcomes, 6. 10.3897/rio.6.e57602.

Walton, S., Livermore, L., Dillen, M., De Smedt, S., Groom, Q., Koivunen, A. & Phillips, S. (2020a). “A cost analysis of transcription systems.” Research Ideas and Outcomes, 6. 10.3897/rio.6.e56211.

Wilson, R.J., De Siqueira, A.F., Brooks, S.J., Price, B.W., Simon, L.M., Van Der Walt, S.J. & Fenberg, P.B. (2022). "Applying computer vision to digitized natural history collections for climate change research: Temperature-size responses in British butterflies." Methods in Ecology and Evolution, 14(2), pp.372–384. 10.1111/2041-210x.13844.

Wu, Y., Kirillov, A., Massa, F. & Girshick, R. (2019). “Detectron2.” GitHub Repository. Available at: https://github.com/facebookresearch/detectron2. [online] [Accessed 05 Jun. 2025].

Zelinsky, A. (2009). “Learning OpenCV—Computer vision with the OpenCV library (Bradski, G.R. et al.; 2008)[On the shelf].” IEEE Robotics & Automation Magazine, 16(3), p.100. 10.1109/mra.2009.933612.

Zooniverse. (2025). “Zooniverse platform.” Available at: https://www.zooniverse.org/. [online] [Accessed 28 Jan. 2025].

