## Supporting Information ELIE for "High Throughput Information Extraction of Printed Specimen Labels from Large-Scale Digitization of Entomological Collections using a Semi-Automated Pipeline" (from another version of this preprint: version 1, 2025-07-12)

### Tables

| Models | Classes | Datasets | Training Images (80%) | Validation Images (20%) | Testing Images (10%) | Total Images | Testing Runtime (sec) | Avg. Runtime Per Image (sec) |
|---|---|---|---|---|---|---|---|---|
| Specimen Identifier Classifier | identifier | Picturae_MfN | 1600 | 400 | 200 | 2200 | 21 | 0.05 |
| | not_identifier | | 1600 | 400 | 200 | 2200 | | |
| Handwritten Printed Classifier | handwritten | Picturae_MfN | 2416 | 604 | 320 | 3340 | 37 | 0.0006 |
| | printed | | 2416 | 604 | 320 | 3340 | | |

**Table 2. Image distribution and runtime benchmarks for the training, validation, and testing of the label classification module. The images were split into 80% training, 20% validation, and 10% testing sets. Results are reported per class across two classification models using the Picturae_MfN dataset. Runtime was measured on an Apple MacBook with an M1 CPU.**

| Model | Classes | Dataset | Training Images (80%) | Validation Images (20%) | Testing Images (10%) | Total Images | Epochs |
|---|---|---|---|---|---|---|---|
| rotation_label | 0 | Picturae_MfN | 272 | 68 | 34 | 374 | 100 |
| | 90 | | 272 | 68 | 34 | 374 | |
| | 180 | | 352 | 88 | 44 | 484 | |
| | 270 | | 352 | 88 | 44 | 484 | |

**Table 3. Distribution of images used for training, validation, and testing in the label rotation module. Images were split into 80% for training, 20% for validation, and 10% for testing. The dataset is organized by rotation angle, with the number of images allocated to each phase of model development shown accordingly.**

| Model | Class | Datasets | Training Images (80%) | Validation Images (20%) | Testing Images (10%) | Total Images | Epochs |
|---|---|---|---|---|---|---|---|
| Label Detection | label | LEPPHIL | 483 | 120 | 60 | 663 | 10 |
| | | Bees & Bytes | 483 | 121 | 60 | 664 | |
| | | AntWeb | 546 | 137 | 69 | 752 | |
| | | Picturae_MfN | 712 | 178 | 89 | 979 | |

**Table 5. Distribution of multi-label images (MLI) used for training, validation, and testing of the label detection module across datasets. The images were split into 80% training, 20% validation, and 10% testing sets. All models used the "label" class, and each dataset was trained for 10 epochs.**

| Models | Input | Training Images (80%) | Validation Images (20%) | Testing Images (10%) | Total Images | Epochs |
|---|---|---|---|---|---|---|
| Specimen Identifier Classifier | Single-label images | 3200 | 800 | 400 | 4400 | 15 |
| Handwritten Printed Classifier | Single-label images | 4832 | 1208 | 640 | 6680 | 32 |

**Table 6. Distribution of single-label images (SLI) used to train, validate, and test the label classification module. Each classifier was trained on a separate dataset, with an 80% training, 20% validation, and 10% testing split among the training, validation, and test sets.**

| Threshold† | Clusters with 1 Label | Clusters with >1 Label | Total Clusters |
|---|---|---|---|
| 0.3 | 46 | 162 | 208 |
| 0.4 | 60 | 195 | 255 |
| 0.5 | 104 | 223 | 327 |
| 0.6 | 179 | 361 | 540 |
| 0.7 | 248 | 403 | 651 |
| 0.8 | 296 | 426 | 722 |
| 0.9 | 609 | 435 | 1044 |

† The minimum similarity score required for items to be grouped into the same cluster.

**Table 10.1. Distribution of clusters in the MfN_LEP_SEASIA dataset at various similarity thresholds. The table shows the number of clusters containing only a single label versus those containing multiple labels.**

| Threshold† | Clusters with 1 Label | Clusters with >1 Label | Total Clusters |
|---|---|---|---|
| 0.3 | 636 | 190 | 826 |
| 0.4 | 735 | 166 | 901 |
| 0.5 | 793 | 150 | 943 |
| 0.6 | 846 | 139 | 985 |
| 0.7 | 892 | 118 | 1010 |
| 0.8 | 1012 | 139 | 1151 |
| 0.9 | 1003 | 235 | 1238 |

† The minimum similarity score required for items to be grouped into the same cluster.

**Table 10.2. Distribution of clusters in the USNM_COL_CAM dataset at various similarity thresholds. The table shows the number of clusters containing only a single label versus those containing multiple labels.**

| Threshold† | Clusters with 1 Label | Clusters with >1 Label | Total Clusters |
|---|---|---|---|
| 0.3 | 431 | 211 | 642 |
| 0.4 | 832 | 169 | 1001 |
| 0.5 | 925 | 168 | 1093 |
| 0.6 | 993 | 180 | 1173 |
| 0.7 | 1150 | 174 | 1324 |
| 0.8 | 1227 | 211 | 1438 |
| 0.9 | 1292 | 227 | 1519 |

† The minimum similarity score required for items to be grouped into the same cluster.

**Table 10.3. Distribution of clusters in the MCZ_ENT_Boston dataset at various similarity thresholds. The table shows the number of clusters containing only a single label versus those containing multiple labels.**

### Figures

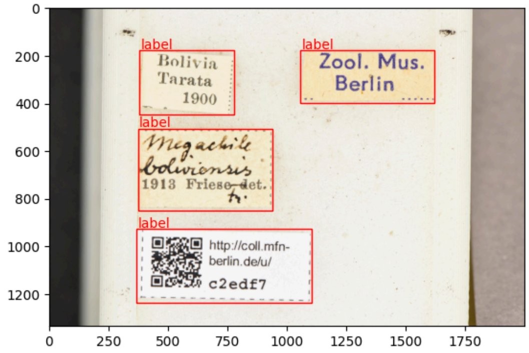


**Figure 3. Example output from the label detection module, displaying bounding boxes and predicted classes ("label") on multi-label images (MLI) before cropping. The boxes were generated by a Faster R-CNN model trained on the LEPPHIL, Bees & Bytes, AntWeb, and Picturae_MfN datasets. A prediction confidence threshold of 0.8 was applied to suppress low-confidence detections. Bounding boxes are shown on uncropped multi-label images from the Bees & Bytes dataset for illustrative purposes (not to scale).**

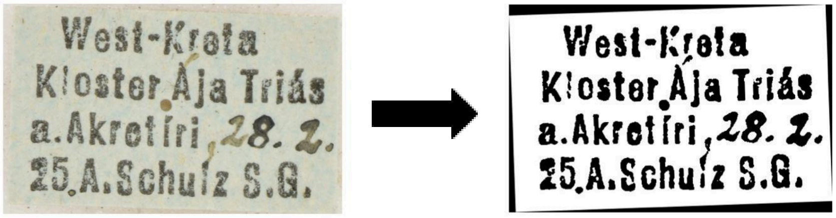


**Figure 5. Example of a single-label image (SLI) from the MfN_LEP_SEASIA dataset before and after image preprocessing. The original image (left) shows uncorrected rotation and background noise, while the processed image (right) has undergone skew correction, noise removal, and contrast enhancement to optimize performance for OCR. This preprocessing step improves text recognition accuracy by standardizing label orientation and improving visual clarity. Images are shown side by side for comparison (not to scale).**

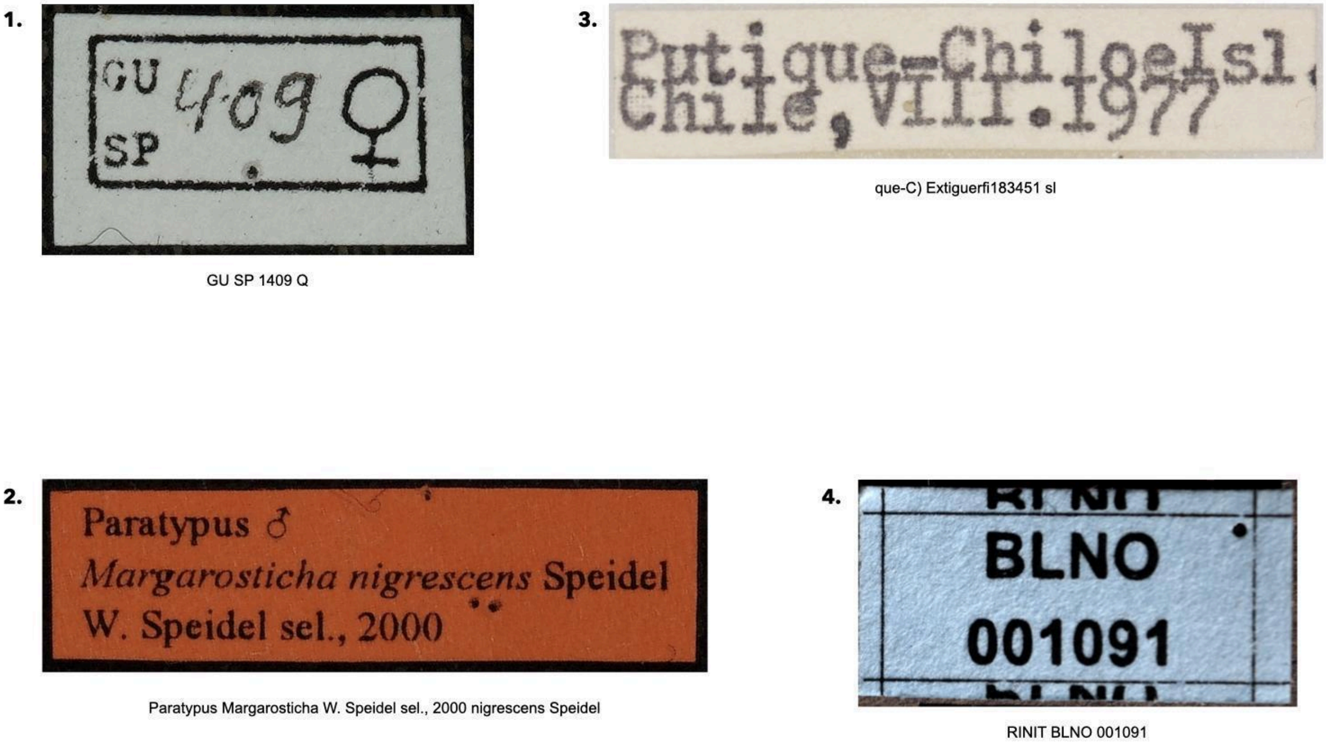


**Figure 13. Examples of common challenges encountered during OCR of insect specimen printed labels, shown alongside their Google Vision API transcriptions. Each panel presents a single-label image (SLI) from the MfN_LEP_SEASIA dataset and its corresponding raw transcription. Specific challenges include uneven ink distribution and inconsistent character spacing (13.1), poor contrast between text and background (13.2), blurred edges, uneven inking, irregular spacing (13.3), and non-standard physical formatting, such as hand-cut labels (13.4). These visual artifacts degrade OCR performance and contribute to increased character and word error rates, even after preprocessing has been applied. Images are not to scale.**
